# A groupwise registration and tractography framework for cardiac myofiber architecture description by diffusion MRI: an application to the ventricular junctions

**DOI:** 10.1101/2021.10.05.463112

**Authors:** Julie Magat, Maxime Yon, Yann Bihan-Poudec, Valéry Ozenne

## Abstract

**Background:** Knowledge of the normal myocardial–myocyte orientation could theoretically allow the definition of relevant quantitative biomarkers in clinical routine to diagnose heart pathologies. A whole heart diffusion tensor template representative of the global myofiber organization over species is therefore crucial for comparisons across populations. In this study, we developed a groupwise registration and tractography framework to resolve the global myofiber arrangement of large mammalian sheep hearts. To demonstrate the potential application of the proposed method, a novel description of sub-regions in the intraventricular septum is presented.

**Methods:** Three explanted sheep (ovine) hearts (size ~12×8×6 cm3, heart weight ~ 150 g) were perfused with contrast agent and fixative and imaged in a 9.4T magnet. A group-wise registration of high-resolution anatomical and diffusion-weighted images were performed to generate anatomical and diffusion tensor templates. Diffusion tensor metrics (eigenvalues, eigenvectors, fractional anisotropy …) were computed to provide a quantitative and spatially-resolved analysis of cardiac microstructure. Then tractography was performed using deterministic and probabilistic algorithms and used for different purposes: i) Visualization of myofiber architecture, ii) Segmentation of sub-area depicting the same fiber organization, iii) Seeding and Tract Editing. Finally, dissection was performed to confirm the existence of macroscopic structures identified in the diffusion tensor template.

**Results:** The template creation takes advantage of high-resolution anatomical and diffusion-weighted images obtained at an isotropic resolution of 150 μm and 600 μm respectively, covering ventricles and atria and providing information on the normal myocardial architecture. The diffusion metric distributions from the template were found close to the one of the individual samples validating the registration procedure. Small new sub-regions exhibiting spatially sharp variations in fiber orientation close to the junctions of the septum and ventricles were identified. Each substructure was defined and represented using streamlines. The existence of a fiber-bundles in the posterior junction was validated by anatomical dissection. A complex structural organization of the anterior junction in comparison to the posterior junction was evidenced by the high-resolution acquisition.

**Conclusions:** A new framework combining cardiac template generation and tractography was applied on the whole sheep heart. The framework can be used for anatomical investigation, characterization of microstructure and visualization of myofiber orientation across samples. Finally, a novel description of the ventricular junction in large mammalian sheep hearts was proposed.

## Background

Mapping cardiac microstructure is essential to understand both the mechanical and electrical heart functions [1, 2]. Although an active field of research, linking the organization of the cardiac microstructure to cardiovascular disease remains challenging. In particular, the mechanisms implicated in triggering and maintaining heart failure [3, 4] or arrhythmias such as persistent AF [5] and ventricular fibrillation [6] are largely unknown.

Several cardiovascular imaging techniques such as micro-computed tomography (μCT) [7, 8] and phase-contrast CT [9–12] combined with structure tensor image (STI) analysis have demonstrated their ability to map the cardiac fiber organization with a spatial resolution close to 10 μm isotropic [13]. They can provide information on myocardial–myocyte (myofiber) orientation and the myolaminar/sheetlet structure orientation. However, *in vivo* translation of phase-contrast X-ray imaging is limited by radiation exposure. Alternatively, destructive techniques like histology can now be performed in three-dimension at 1 μm isotropic in-plane resolution [14] and allow the assessment of cell-type distribution.

Cardiovascular diffusion tensor imaging (DTI) is a non-destructive method, allowing measurement of myocardial microstructure both *ex vivo* [15, 16] and *in vivo* on humans [17] or animals models [18]. The clinical application of cardiac DTI was recently highlighted by several studies demonstrating how the variation of the microstructure between systole and diastole can be a marker of dilated cardiomyopathy [19] and hypertrophic cardiomyopathy [20]. Other *in vivo* studies [21, 22] have also demonstrated that changes in diffusion tensor metrics, such as Fractional Anisotropy (FA), and Apparent Diffusion Coefficient (ADC) agree with late gadolinium enhancement (LGE) imaging and can depict areas with myocardial infarction and tissue remodeling. Despite growing progress in the field of MRI pulse sequences [23–26], hardware [27] and reconstruction methods [28], clinical *in vivo* DTI MRI remains limited in spatial resolution and signal to noise ratio (SNR) by clinical field strength, and scan times [29]. *In vivo* cardiovascular DTI must also take into account combined respiratory and cardiac motions, and compensated for them with a dedicated acquisition scheme [23]. In contrast, *ex vivo* cardiovascular DTI acquisition at ultra-high field on a static cardiac sample simplifies the acquisition scheme and allows for enhanced SNR, spatial and/or angular resolution. It therefore offers a better image quality compared to clinical acquisition providing an excellent research tool for anatomical investigation or validation of *in vivo* techniques.

Among all the *ex-vivo* studies using DTI, there is a large amount of evidence depicting the typical organization of the left ventricle (LV) myocardium characterized by the helix angle variation through the transmural wall [28–31]. Only a few studied describe sheetlets arrangement and fiber organization in both the LV and RV [30, 31]. However, in our knowledge, there is a lack of a recent and high-resolution description of the microstructure organization of the junctions between the free wall of the left and right ventricles and the intraventricular septum (IVS) in large mammalian models. A few decades ago, Kuribayashi and Robert [32] demonstrated the existence of abnormality of the myocardium at the junction associated with hypertrophic cardiomyopathy. By macroscopic inspection of tissue sections, the authors classified the histological characteristics of these junctions according to “myocardial disarray”. Therefore, knowledge of normal fibers organization could be valuable for the identification of myocardial disarray and risk stratification [32].

The gold-standard method in image processing for creating a baseline map sharing spatial information with a group of subjects/samples (for instance a structural map describing the anatomy or the fiber organization and an associated parcellation) is named atlas or template. Such a framework normalizes the individual data to a common coordinate space and promotes the comparison between subjects of qualitative and quantitative information with one or multiple imaging modality sharing (or not) the same level of resolution. Spatial normalization is widely used in neuroimaging studies for structural and functional studies [33–35]. In cardiac microstructure studies, *ex vivo* DTI atlas has been initially proposed in 2006 on canine heart (0.3 × 0.3 × 0.9 mm^3^) [36] and was applied later in human (2× 2× 2 mm^3^) [37, 38], pig [39], murine (0.043 × 0.043 × 0.043 mm^3^) [40]. However, except for the murine study, these atlases have been created with low or anisotropic spatial resolution diffusion weighted (DW) images, and suffer from important voxel averaging of diffusion parameters.

In this work, we develop a template-based and tractography framework allowing us to resolve cardiac microstructural fiber anatomy of sheep hearts. An atlas is created with 3 *ex vivo* female sheep hearts fixed with high robustness sample preparation and acquisition protocol [16]. The template creation takes advantage of anatomical and diffusion-weighted images acquired at 9.4 T at an isotropic resolution of 150 μm, 600 μm respectively providing an accurate region-based analysis of myofiber architecture. We also emphasize the use of tractography to depict the global myofiber organization. To demonstrate the potential application of the proposed atlas and tractography visualization, we provide a novel description of sub-regions in the IVS and LV in which fibers present either an abrupt change in orientation or a notable deviation from the circular arrangement of the fibers depicted by the helix angle rule. The study focuses on: i) the presence of a singularity/fascicle of fibers at the junction of the posterior wall while noticing that the fiber arrangement mostly continues between both RV and LV wall and the septum. ii) the complex organization of the anterior junction where a division of the septal wall into the two fiber-bundles going through the aorta and pulmonary artery is visible iii) a singularity/fascicle close to the anterior junction going to the papillary muscle.

## Methods

### Sample Preparation

The hearts of 4 female sheep (ovine) were explanted via sternal thoracotomy under general anesthesia, This protocol was approved by the Animal Research Ethics Committee in accordance with the European rules for animal experimentation. Three hearts were dedicated to MR experiments and one heart was dissected for macroscopic visualization. The three hearts (~12×8×6 cm^3^, heart weight=150±10 g) were in relaxed state due to a first cardioplegic flushing before formalin fixation. They were then perfusion-fixed in 10% formaldehyde containing 2 mL of gadoterate meglumine, a gadolinium-based contrast agent to enhance the contrast between the myolaminae and cleavage planes and decrease the longitudinal relaxation time. To attenuates susceptibility artifacts at the border of the cardiac chamber, the heart were removed from the solution and the cavity were filled with Fomblin, a perfuoropolyether with no ^1^H signal when scanned using MRI. Lastly, whole hearts were immersed in Fomblin and sealed in a plastic container without rehydratation. This procedure is described with additional details in Magat et al. [16].

### *Ex vivo* anatomical and DWI acquisition

All MRI acquisitions have also been described previously in [16]. A summary is as follows: the experiments were performed at 9.4T (Bruker BioSpin MRI system, Ettlingen Germany) with an open bore access of 30 cm and a 200-mm inner diameter gradient (300 mT/m).

- A 3D FLASH sequence was applied to get anatomical images of the whole heart volume, at an isotropic resolution of 150 μm. The total acquisition time for each heart was 30 h 50 min. The resulting volume is referred to as the anatomical image. The parameters were : TE = 9 ms, TR = 30 ms, matrix size = 731 × 665 × 532, FOV = 110 × 100 × 80 mm^3^, flip angle = 30°. An acceleration factor in the first phase dimension of 1.91 was used and the images where reconstructed with GRAPPA. The total acquisition time for each heart was 30 h 50 min.
- Diffusion-weighted (DW) images were acquired using a 3D diffusion-weighted spin-echo sequence at an isotropic resolution of 600 μm with 6 diffusion encoded directions, a single b-value of 1000 s/mm^2^ and one b0 image. The parameters were : TE = 22 ms, TR = 500 ms, FOV = 100 × 80 × 110 mm3, matrix = 166 × 133 × 183. An accelerator factor in phase direction of 1.8 was used and the images where reconstructed with GRAPPA. The total acquisition time for each heart was 16 h 55 min.

Each dataset was reconstructed with Paravision 6.0 software (Bruker, Ettlingen, Germany)

### Anatomical and DW images processing for template generation

The steps performed to generate the anatomical and the diffusion tensor template are detailed below and summarized in Fig. 1. Unless otherwise mentioned, all registration steps and template generation were performed with ANTs library with the default parameters [41]. Estimation of diffusion tensor and derived tensor maps were performed using the MRtrix3 software [42, 43]. All the processing were fully automatic, reproducible and executed using shell scripts without any input from the user with the exception of long axis alignement and ROIs and seeds definition. The corresponding transformations, ROIs and seed have been shared for reproducibility purpose (see data avaibility section).

**Figure 1.**
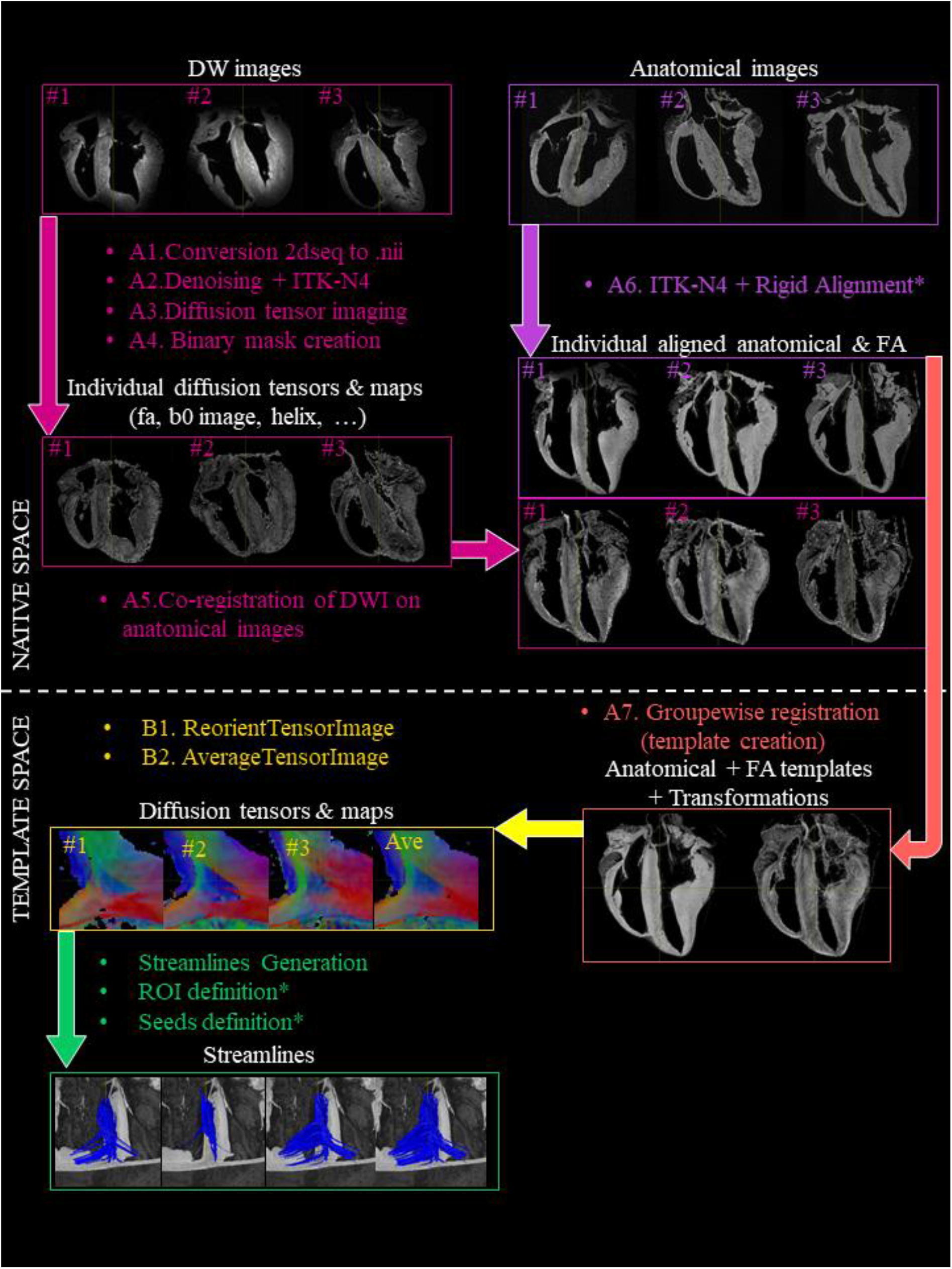
block diagram of the processing steps. A) Anatomical and DW images were first handled separately with a B1 inhomogeneity field correction. Then, anatomical images were manually aligned along the long axis using ITK-snap while diffusion tensor calculations were performed before any registration. Secondly, DW images were registered to the anatomical images. Model generation was performed using the symmetric normalization transformation (SyN) model using the anatomical and FA images. B) The individual diffusion tensors were then deformed into the template space and an average diffusion tensor was created. After this step, the diffusion tensors and maps and streamlines are spatially normalized and available individually or jointly (averaged). Manual steps are indicated (*), all other steps are fully automatic, reproducible and executed using shell scripts.

#### A) Anatomical and FA template generation

1. Conversion from Bruker to Nifti format. Anatomical and DW images 2dseq files were converted in Nifti format with the dicomifier library (https://dicomifier.readthedocs.io/).
2. DW images pre-processing. Each of the DW images series was up-sampled by a factor of 2 using trilinear interpolation to reach a voxel size of 0.3×0.3×0.3 mm^3^. A denoising was performed using the MP-PCA method [44]. Finally, a B1 inhomogeneity field correction was applied with ITK-N4 [45].
3. Diffusion tensor imaging estimation. Diffusion tensor calculations were performed prior any registration to avoid difficult transformation of the diffusion encoding matrix. Diffusion tensor maps (eigenvalues: λ_1_, λ_2_, λ_3_, ADC, FA, and color-coded FA (cFA) also known as Red-Green-Blue colormap) were obtained with MRtrix3. The first diffusion tensor eigenvector v1 corresponds to myofiber’s main orientation [46, 47]. The second v2 and third v3 eigenvectors were associated with sheetlet in-plane and normal directions, respectively.
4. Estimation of the cardiac mask. A binary mask was created and applied on both anatomical and DW images to segment myocardial tissue from regions with hypersintense signal corresponding to residual formaldehyde. Low and high cutoff thresholds were applied on the FA, trace, and DW images to define the binary mask as described previously in [16].
5. Co-registration of DW images on anatomical images. Anatomical and DW images were not always acquired during the same session because of the long scan time and scanner availability. Therefore, alignment of each DW image on its respective anatomical images was needed and performed using Affine Registration using ANTs. A B1 homogeneity using ITK-N4 was also performed on anatomical images before the co-registration. Finally, FA maps were mapped to the space of the anatomical image as shown in Fig. 1A.
6. Rigid Alignment of the anatomical and FA images. Images of each sample (Anatomical and FA) were registered manually with ITK-snap [48] to align the long axis of the LV to the Z-axis of each volume and hence, obtain a similar alignment among hearts.
7. Template generation. The template generation [33], from coarse resolution to the final resolution, was performed iteratively with 4 levels of resolution (1 mm, 0.6 mm, 0.3 mm, 0.15 mm). Anatomical and FA images were used jointly using the following modality weights 1 × 0.5 respectively.
  - At each level, the templates generated from the previous level were resampled to the current resolution and used as input to guide the current template creation using the standard script ‘antsMultivariateTemplateConstruction2.sh’ with the symmetric normalization transformation model (SyN) [49].
  - For the first level (1 mm), the initialization sequence of template creation was similar to the one published in [50].

The same iterative strategy was performed to generate additional templates using only rigid transformation and affine transformation for comparison (Fig. 2–3). After this step, the set of forward (or inverse) transformations for moving from individual subject’s space (also known as native space) to template space (or the inverse) were obtained. To determine registration quality, the Dice Similarity Coefficient [51]] and Jaccard index [52] between each individual cardiac mask were computed for different transformation models (without transformation, after manual alignement, after Rigid, Affine and SyN registration).

**Figure 2:**
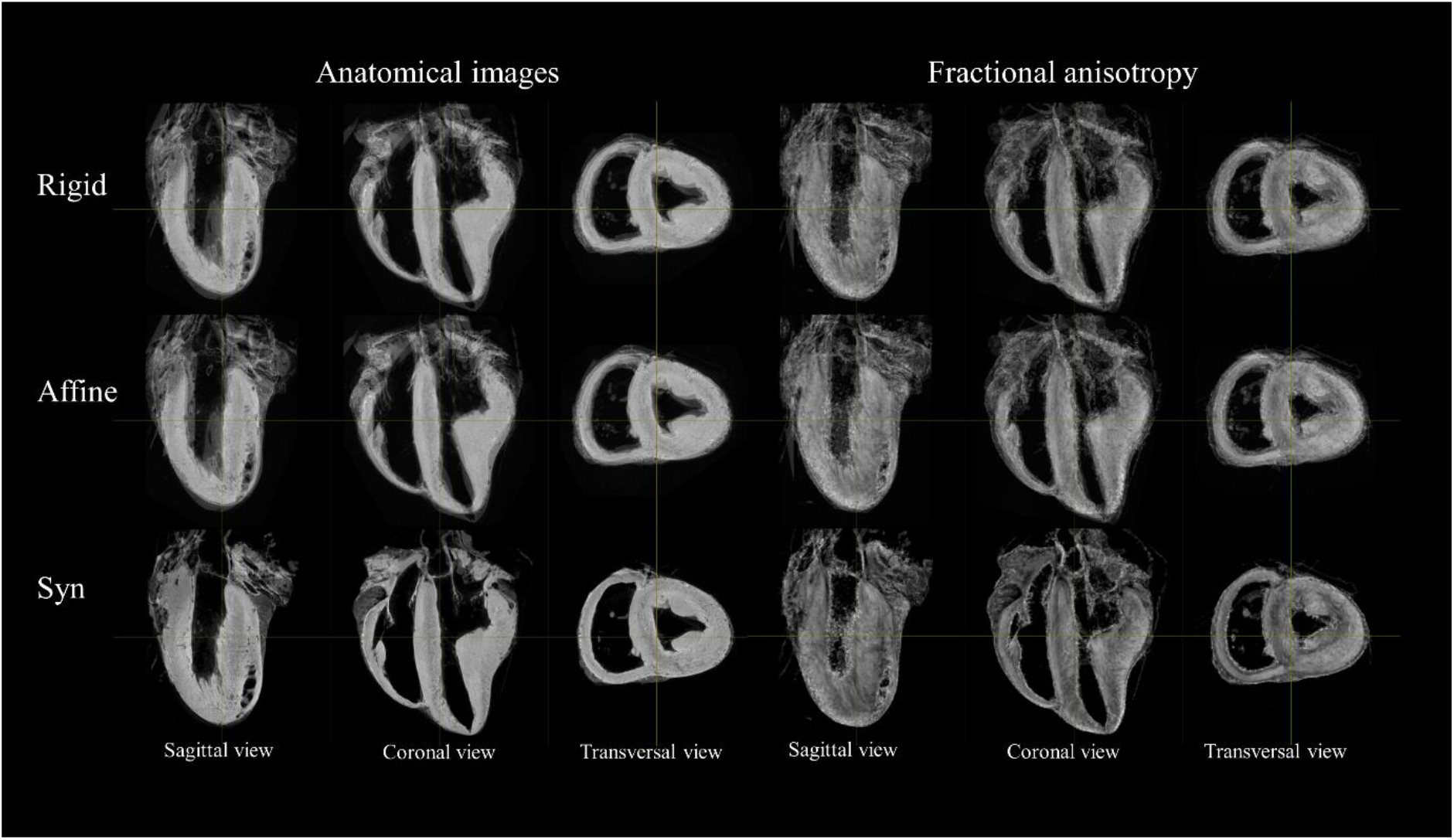
Impact of the transformation model during the registration on anatomical and FA template. Sagittal coronal and transverse view of anatomical (left) and FA (right) template using Rigid (top row), Affine (middle Row), SyN (bottom row). Mis-registration areas using Rigid or Affine transform are visible in the basal part of the sample, at the apex or on the papillary muscle.

**Figure 3:**
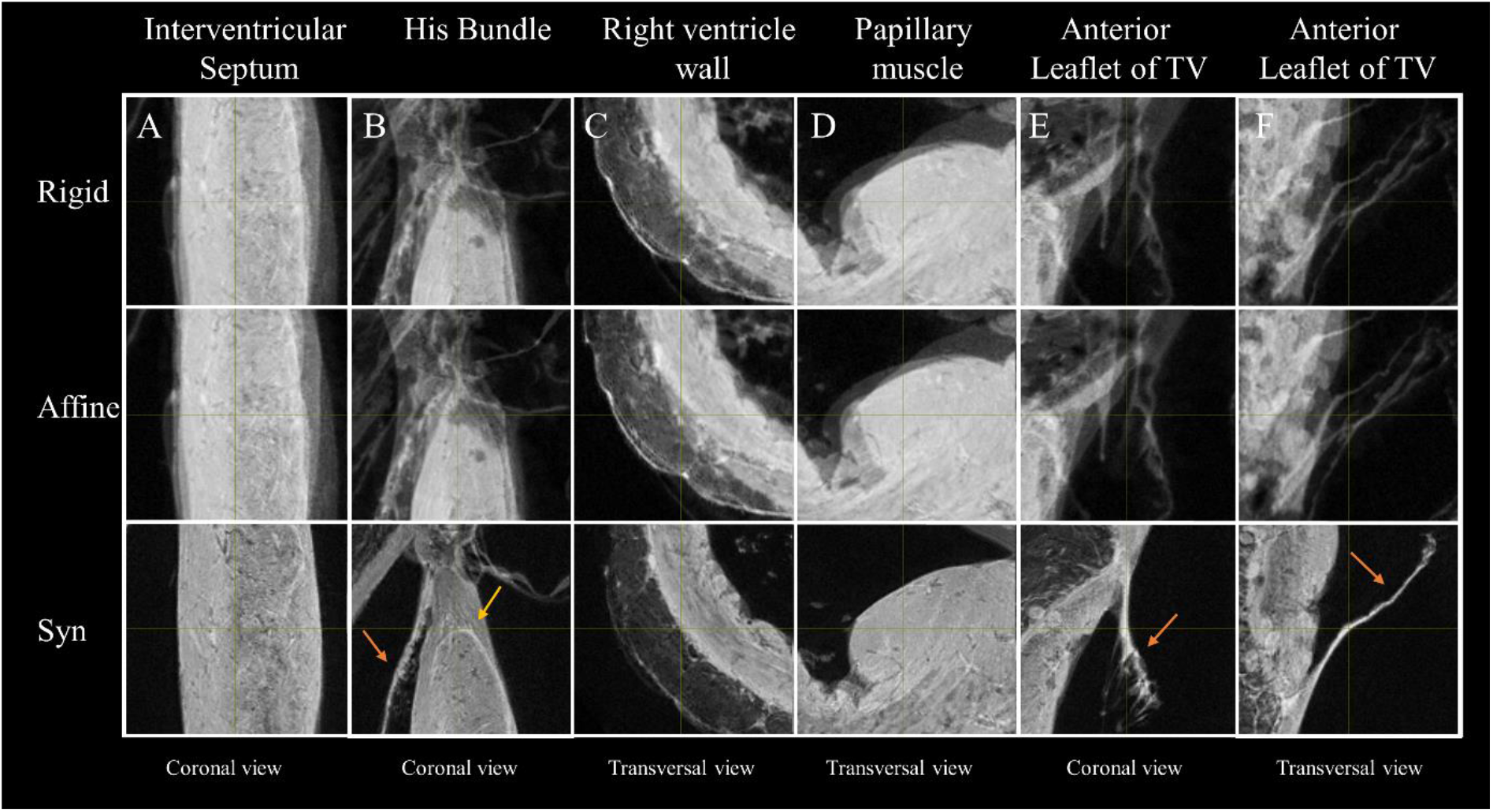
Impact of the transformation model during the registration on the anatomical template in five different regions. Zoom-view in the IVS (A), near the division of the His bundle in left and right bundles (B), in the RV wall (C), in the posterior papillary muscle of the LV (D) and on the anterior leaflet of the tricuspid valve (TV) (E and F) using Rigid (top row), Affine (middle Row), SyN (bottom row) transformation model, The misregistration areas visible using Rigid and Affine transformation are less pronounced using SyN for the large or small structures like the conduction system (yellow arrow) or the leaflets of the valve (orange arrows).

#### B) Diffusion tensor template generation

1. Mapping of individual diffusion tensor images to the template space. Individual diffusion tensor images were warped in the template space. This process includes both registration of the voxel at the new location and reorientation of the tensor via the ‘antsRegistration’ and ‘ReorientTensorImage’ command of ANTs. Preservation of eigenvector orientations related to the reorientation of the sample from the native space to the template space was carefully checked during all processing steps.
2. Diffusion tensor template generation and computation of derived maps. The transformed diffusion tensors were averaged via the AverageTensorImages’ of ANTs to create the diffusion tensor template. Diffusion tensor maps were computed as described in the previous section A.3.

### Tractography processing

Tractography was used for different purposes: i) Visualization of fibers organization, ii) Segmentation of sub-area depicting the same fiber organization, iii) Seeding and Tract Editing. All the tractography steps described in this section were performed with MRtrix3 [42]. For each application, different algorithms were tested:

- The deterministic tracking algorithm FACT [53] where streamline trajectory is determined by the principal eigenvector of the diffusion tensor.
- Tensor_Det [54], a deterministic approach with intra-voxel interpolation denoted here as DET.
- Tensor_Prob [55], a probabilistic algorithm with intra-voxel interpolation denoted here as PROB.

Unless specified, we defined the tractography parameters as an FA-stopping threshold of 0.1, a maximum angle between steps of 60 degrees, a step size of 0.05 mm, a maximum length of 40 mm, a minimum length value of 1 mm.

Streamlines generated by FACT can be computed in the native space using the primary eigenvector of the individual diffusion tensors or in the template space using the primary eigenvector of the average diffusion tensor. The average diffusion tensor is depicting an averaged myofiber organization across samples. Streamlines generated by DET and PROB needed to be computed in the native space and warped in the template space to ease the comparison. Seeding ROI and tract filtering region of exclusion/inclusion were manually drawn in the template space and warped to the native space.

Tractography was performed on the posterior singularity with the three mentioned algorithms with varying parameters: max. length: 10 mm, 20 mm, 40 mm, 60 mm, angle: 15°, 30°, 45°, 60°, FA-stopping threshold 0.1, 0.15, 0.2, 0.25, step size 0.01 mm, 0.05 mm, 0.1 mm, 0.2 mm.

In the anterior and posterior junction, four cylindrical seeding regions (R1, R2, R3, R4) were manually drawn at the LV/RV junction in the apex, mid-ventricular and base area. Based on the location of the region of interest (ROI) three pathways were arbitrarily defined to study the fiber organization at the junction: pathway A (from ROI R1 to R2) was the pathway inside the free wall of the RV. Pathway B (from ROI R1 to R2 to R3) described a pathway from the RV to the beginning of the free wall of the LV while pathway C (from ROI R1 to R2 to R4) described a pathway from the RV to the IVS. For each pathway, streamlines were computed and filtered by identifying tracts that pass through all the corresponding ROIs. For each tracking algorithm tested, the resulting number of streamlines were counted.

### Dissection study and macroscopic anatomy visualization

The heart dedicated to macroscopic examination was dissected as follows.

- The LV, RV and then the IVS were transected from base to apex to remove the anterior part of the heart and isolate the posterior wall of the heart.
- Transverse (short-axis) cuts of the sample were performed progressively from base-ventricular level to mid-ventricular level.
- A coronal cut (long axis) was performed on line ranging from the RV cavities to the LV posterior wall going through the suspected region.

Photos were taken with a Canon PowerShot SX710 HS and a Samsung Galaxy SM-G960F with the surgical light of the operating room. Rigid registration was manually performed using ITK-snap to align photography on anatomical images.

## Results

Figure 2 shows sagittal, coronal and transverse views of the anatomical and FA templates with different transformation models used during the template creation. The template includes not only the ventricles but also the basal parts of the heart, including the valve, atria, the aorta or the pulmonary artery. Misregistration areas were more visible with rigid or affine transformations than with the SyN transformation model. Several differences are evident through visual inspection: (i) a sharper definition of the myocardial wall (Fig. 2, Fig. 3A-F); (ii) an enhanced definition of the division of the His bundle in left and right bundles (Fig. 3B); (iii) a more accurate delineation of the leaflets of the tricuspid valve (Fig. 3B, 3E, 3F). Consistently misregistered areas using the SyN transformation model were noticeable in the basal part (Fig. 2) and for most of the small or large vessels (Fig. 3E) going through the myocardium or around (like the coronary sinus) due to the variability of the vessel locations.

Supplementary Fig 1A and 1B shows the Jaccard (A) and Dice (B) metrics computed between each individual cardiac mask for different transformation model. Using quantitative image-based metrics, the Symmetric Normalization (SyN) transformation model showed superior performance over other transformation models and were verified by pariwise T-tests in Supplementary Fig 1C. Supplementary Fig. 2A shows the diffusion metric distributions for the three individual diffusion tensors and the averaged diffusion tensor. All the voxels of the left ventricle are included. The mean FA is at 0.28 ± 0.08, 0.25 ± 0.07, 0.27 ± 0.9 and 0.25 ± 0.06 for sample #1, #2, #3 and averaged diffusion tensor respectively. The mean ADC values for sample #1, #2, #3 and averaged tensor were 8.76 ± 1.32, 9.32 ± 1.21, 9.37 ± 1.53 and 8.76 ± 1.32 (x10^−4^) mm^2^/s respectively. The mean first eigenvalues (λ1) computed values for sample #1, #2, #3 and averaged tensor were = 11.43 ± 1.28, 11.88 ± 1.2, 12.1 ± 1.45 and 11.38 ± 0.97 (×10^−4^) mm^2^/s.

cFA maps overlaid on anatomical data are provided in five Supplementary movies 3A-E (for each sample #1, #2, #3, on the average diffusion tensor images, and all together, respectively). All datasets were warped in a template space to ease the comparison. In the short-axis view, the blue color indicates that fiber orientation in the papillary muscle or in the IVS is parallel to the long axis on the heart. On the other hand, green and red colors indicate the orientation of the fibers in the short axis plane in Anterior-Posterior (AP) or Left-Right (LR) orientation respectively. While a smooth gradient of color from green to red (or vice versa) indicates a normal arrangement of the cardiac myofibers, the existence of small sub-regions in the LV indicates either a rapid change in orientation or apparent boundaries. Especially: fiber arrangements in Inferior-Superior (IS) orientation are visible: i) from the basal area to the mid-ventricular level close to the papillary muscle, ii) at the intersection of the LV and RV from the basal area to mid-ventricular level, iii) in the basal area of the IVS, divided into two fiber-bundles that go on the one hand to the pulmonary artery and on the other hand to the aorta.

To further investigate the existence of abrupt changes in orientation, we first focused our analysis on the posterior wall. Fig. 4A-C displays coronal, sagittal and transverse views of the anatomical and FA templates at the junction of the RV, LV and IVS in the basal area. Fiber orientation can be visually identified on the anatomical template images due to 150 μm resolution and the T2* contrast. A brighter image intensity is noticeable in the layer of the IVS close to the RV and at the junction of the ventricles. The FA derived from the diffusion tensor template is relatively homogeneous except for two regions with lower values. They are i) a line dividing the IVS into two areas and ii) a triangle located at the junction. Fig. 4D and E displays the cFA maps and the streamlines respectively with the anatomical image as background. Smooth transitions of fiber orientation are visible (color gradient of green to purple) in the endocardial part of the LV while fiber orientation changes abruptly between adjacent voxels in the IVS and in the middle of the LV/RV junction (blue) depicting a triangular shape in the coronal view. The triangular shape will be denoted as the posterior singularity in the rest of the article. Note the presence of a channel or fascicles of fibers going from the RV to the IVS (in green).

**Figure 4:**
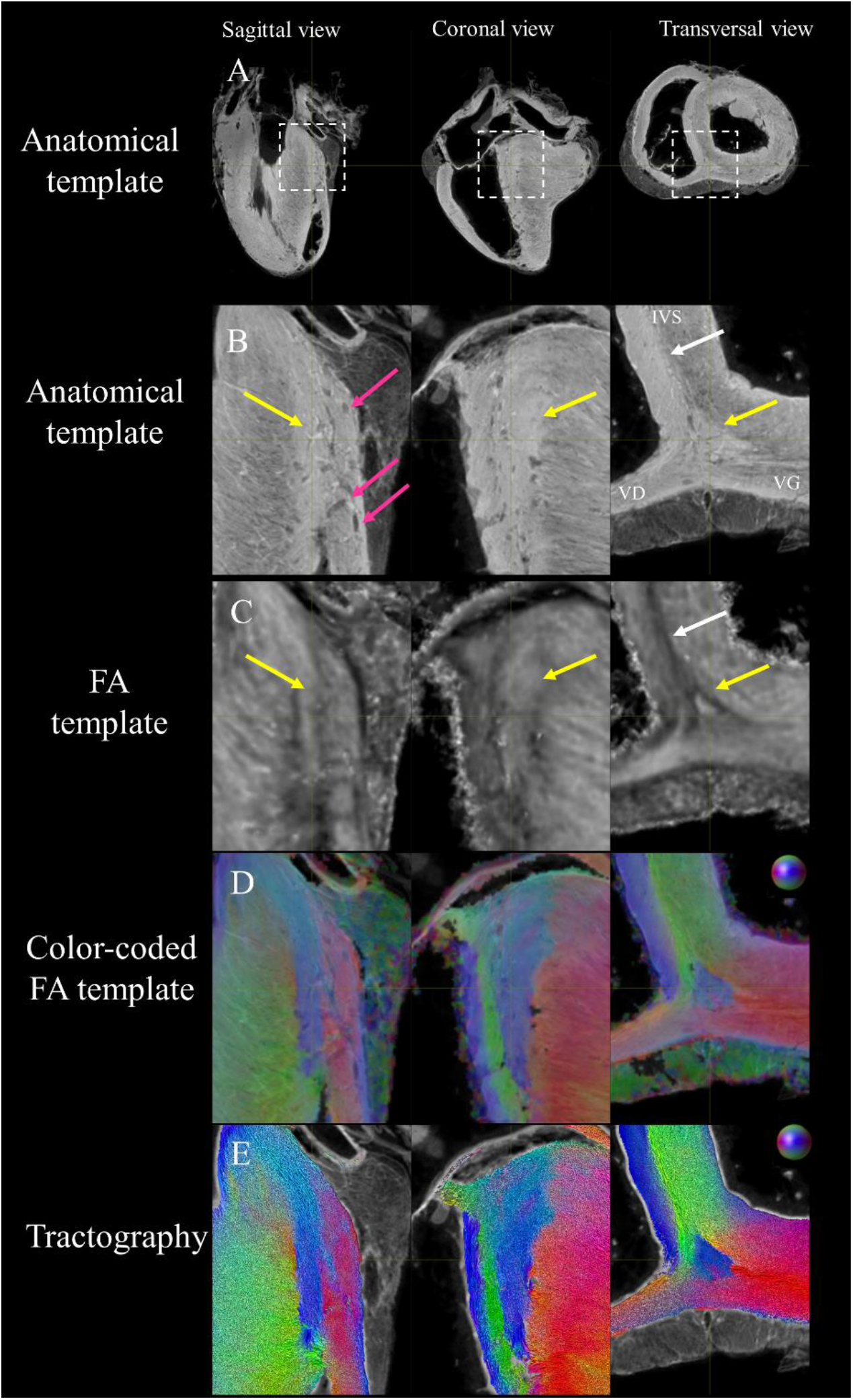
Visualization of the posterior LV/RV junction in the basal area using anatomical, FA and diffusion tensor templates in sagittal, coronal and transverse view. (A) Large view and (B) zoom-view of the anatomical template on the posterior LV/RV junction in the basal area. Myocardium with different grey levels and epicardial fat are visible. (C) FA template. Lower fractional anisotropy value is noticeable. (D) cFA maps overlaid on anatomical data. Smooth transition of fiber orientation is visible (gradient of green to purple) in the endocardial part of the LV while fiber orientation changes abruptly between adjacent voxels in the IVS and in the middle of the LV/RV junction (yellow arrow) depicting a triangular shape in the coronal view. (E) streamlines with the color-coding of the cFA maps (D). Note the presence of a channel or fascicles of streamlines going from the RV to the IVS (in green).

Fig. 5A and 5B display a coronal and sagittal view of the heart respectively. To give an overview of the size of the fiber arrangement, a masking region of interest was semi-automatically defined to display the posterior singularity independently of the surrounding structures. As a consequence, streamlines leaving the ROI were cut in the representation. It represents a volume of 1135 mm^3^ and a surface of 845 mm^2^. Different tracking techniques and representations are shown for each individual diffusion tensor in Fig. 5C-E. Here, few seeds are manually defined, then fiber tractography streamlines are computed without any ROI restriction. The resulting fiber arrangement of each individual sample presents a base-apex orientation and is progressively continuing within the RV and LV. Supplementary Fig. 4 shows the tractograms obtained with different tuning parameters and the three tracking algorithms (FACT, DET, PROB). Streamline termination was defined either by the cutoff parameter or forced by the max length parameter. A maximum angle defined between 15 and 60° resulted in plausible streamlines. All representations confirm the apex-base orientation of the posterior singularity while the parameters influence the potential connectivity with the surrounding structure.

**Figure 5:**
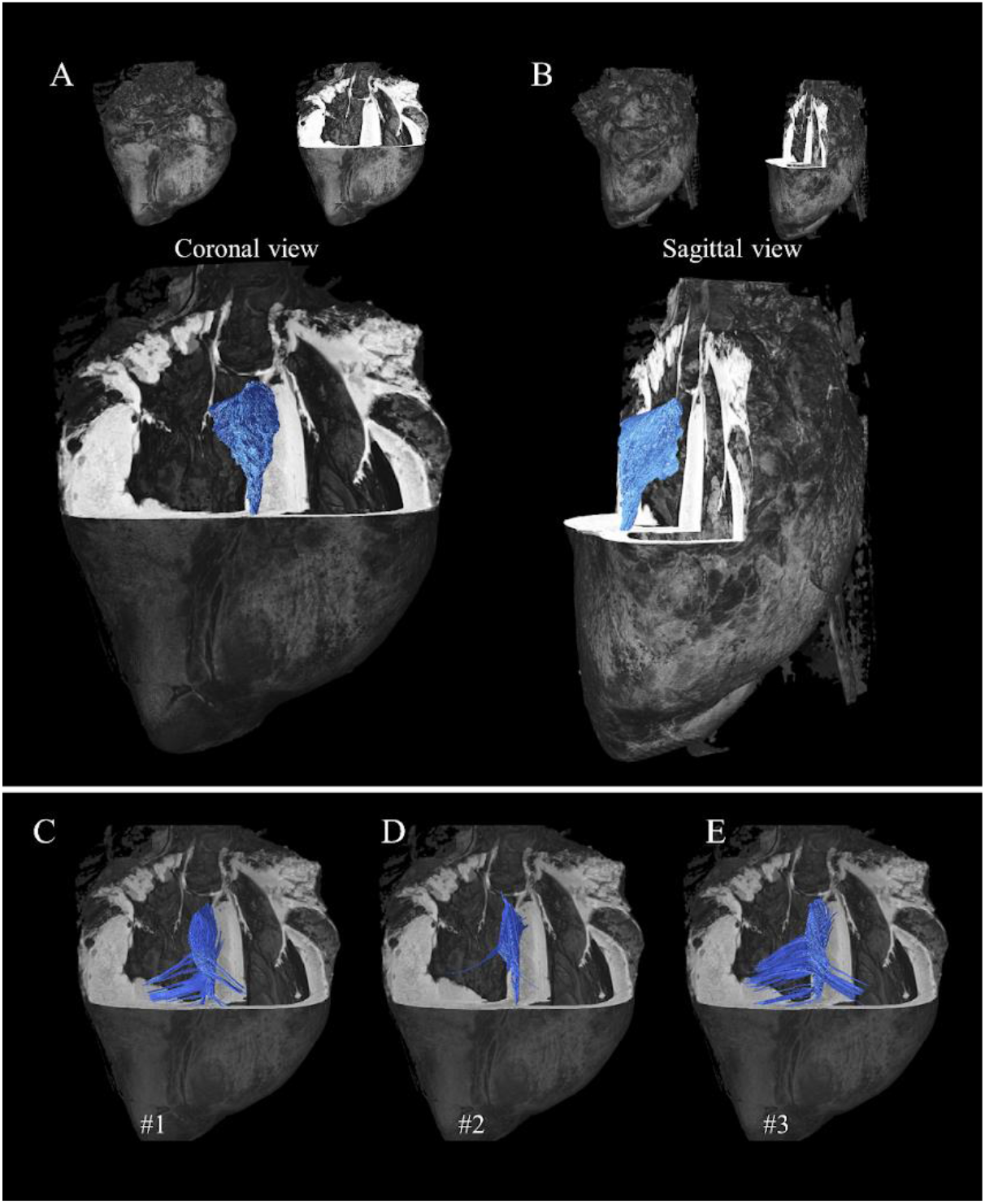
Streamlines of the posterior singularity with different tracking techniques on template and individual diffusion tensors. For each representation, the anatomical template is inserted for reference. Top) Streamlines were computed using a masking region of interest and the FACT algorithm on the primary eigenvector of the average tensor in template space and displayed in coronal (A) and sagittal view (B). Down) A different tracking technique is shown. Few seeds were manually defined, then streamlines were generated for sample #1 (C), #2 (D), #3 (E) using the Tensor_Det algorithm on each individual tensor in native space and warped into template space without ROI restriction.

Anatomical images of the IVS and the picture of the dissected sample are shown in transversal orientation in Fig. 6A and Fig. 6B, respectively. An overlay of both images is shown in Fig. 6C after rigid registration and shows an acceptable agreement by visual inspection. The coronal cut in the posterior area is indicated by a yellow line ranging from the RV cavities to the LV in Fig. 6D. A section of the dissected sample is presented using five pictures for which the sample has been slightly rotated (Fig. 6E-I). Images reveal macroscopic features depicting the fiber orientation changes and support the evidence of edges. As a comparison, streamlines are displayed in short axis and long axis orientation in Fig. 6J and 6K respectively. The fiber orientation display between Fig. 6K and Fig. 6G or 6L shows similar fiber orientation.

**Figure 6:**
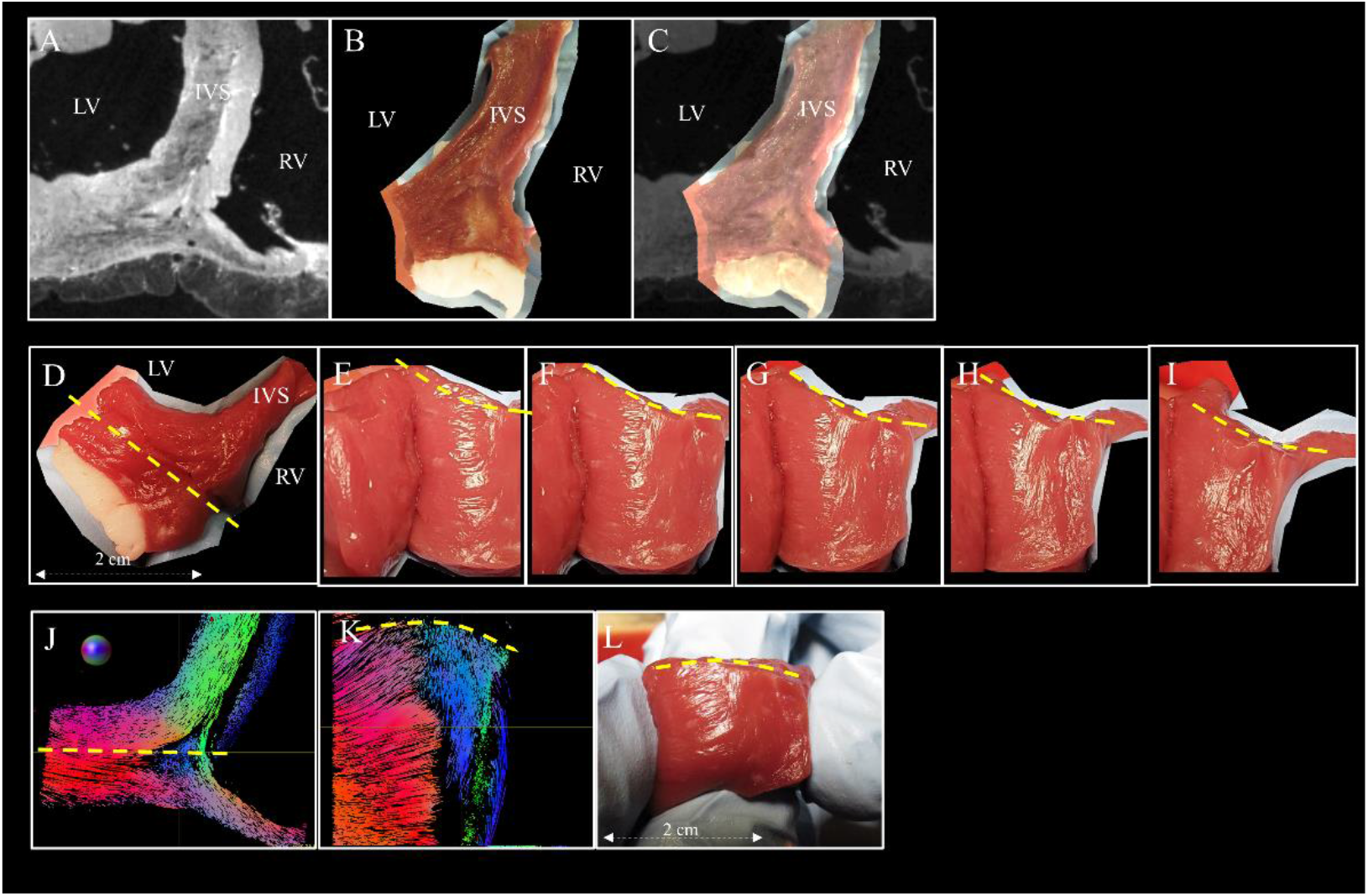
Macroscopic structural organization of the posterior junction. A) Anatomical Transversal image of sheep heart #2 of the posterior junction closed to the base-ventricular level. B) Photography of the dissected heart #4. Corresponding view using anatomical images from sample #2 after rigid registration. C) Overlay of A) over B). D) Another transversal view of the posterior junction. A coronal cut was performed on a line (yellow line) ranging from the RV cavities to the LV posterior wall going through the suspected region. E to I) Visualization of the structural organization is presented by slightly rotating the sample while the surgical light highlights the macroscopic structural organization. J) Tractography in short axis view taken from sample #2 in a selected slice in the basal area K) Tractography in a long-axis view corresponding to the plane drawn by the yellow line. L) Another picture was taken in a long-axis view. Original photo are available to the reviewers.

Fig. 7A-B displays the cFA maps of the anterior junction close to the base in axial and sagittal views. The white dash line indicates the plane of the sagittal view. A smooth transition of fiber orientation is visible in the LV and the endocardium part of the IVS (with a gradient of green to purple). In the epicardium part of the IVS, we observed a division of the fibers inside the IVS into the fiber-bundles going through the aorta (yellow arrow) and the pulmonary artery (purple arrow). The third fascicle of fibers (in pink) located close to the endocardium in the LV was also noticeable with an orientation differing from the surrounding structures. For each mentioned area, streamlines were computed using seeds on the averaged diffusion tensor with the FACT algorithm. To facilitate the rendering, the sample is flipped and the wall of the pulmonary artery and the aorta is manually segmented and plotted as a surface mesh in Fig. 7C-H. Tractography shows a continuity between the IVS and the wall of the pulmonary artery (in green Fig. 7F, I), a continuity between the IVS and the wall of the aorta (in green Fig. 7G, I) and the existence of fibers that split into two branches: one going to the papillary muscle, the other surrounding the top part of the LV (in red Fig. 7H,I). Some streamlines going from PA to the LV and from AO to the RV (yellow arrows) are also found close to the epicardium of the ventricles.

**Figure 7:**
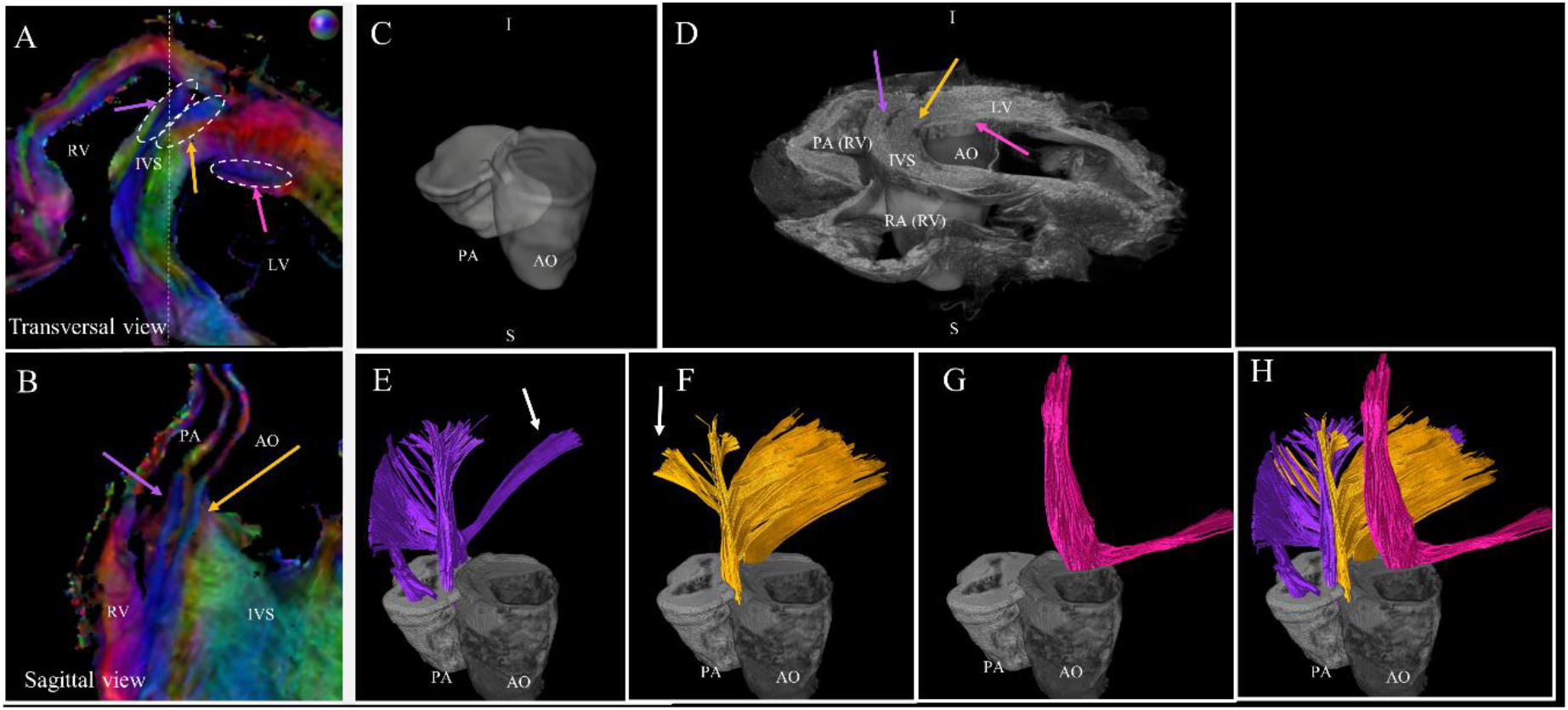
Tractography of the anterior junction. A) and B) zoom-view of the cFA maps of sample #1 in the anterior junction in the basal area in transversal and sagittal orientation, respectively. Three regions of interest are identified: a fascicle of streamlines coming from the IVS splits into two branches going on one side to the wall of the pulmonary artery (purple arrow) and, on the other side to the wall of the aorta (yellow arrow). The third fascicle of streamlines, located in the LV, goes from the aorta to the papillary muscles (pink arrow). C & D) 3D rendered image of the anatomical template with the superimposed mesh of the PA (white) and the AO (grey) Note that the volume was flipped and this orientation was kept for all following representations. E,F,G,H) For each region, seeds were manually defined, then streamlines were computed using the template diffusion tensor without ROI restriction.

To illustrate the added value of the seed-based tractography and analyze the global myofiber architecture, three arbitrary pathways of fibers are defined in the anterior and posterior junctions in the mid-ventricular area. The resulting streamlines for sample #1 are proposed in Fig. 8 in 3D rendering views. In the posterior wall (down), the connection of fibers originating from the RV into the LV free wall and the IVS is detectable (note that the unidirectional direction is arbitrary) (pathway A). Existing pathways (B) and (C) were found connecting. In the anterior wall (top), streamlines split into two output tracks but the fascicle of fibers in the IVS leaves the image plane and goes to the wall of the pulmonary artery as described in Fig. 7. Existing pathways (B) and (C) are found but with a predominant direction toward the free wall. The corresponding results in the basal area are shown in Supplementary Fig. 4. In the posterior wall (down), streamlines split into multiple output tracks located either in the IVS, in the junction or the free wall (pathway A). Existing pathways (B) and (C) are found connecting. On the contrary, in the anterior wall (top), the connection of fiber originating from the RV into the LV free wall is limited to output tracks located on the epicardium surface of the sample and the pathway (C) is not found.

**Figure 8:**
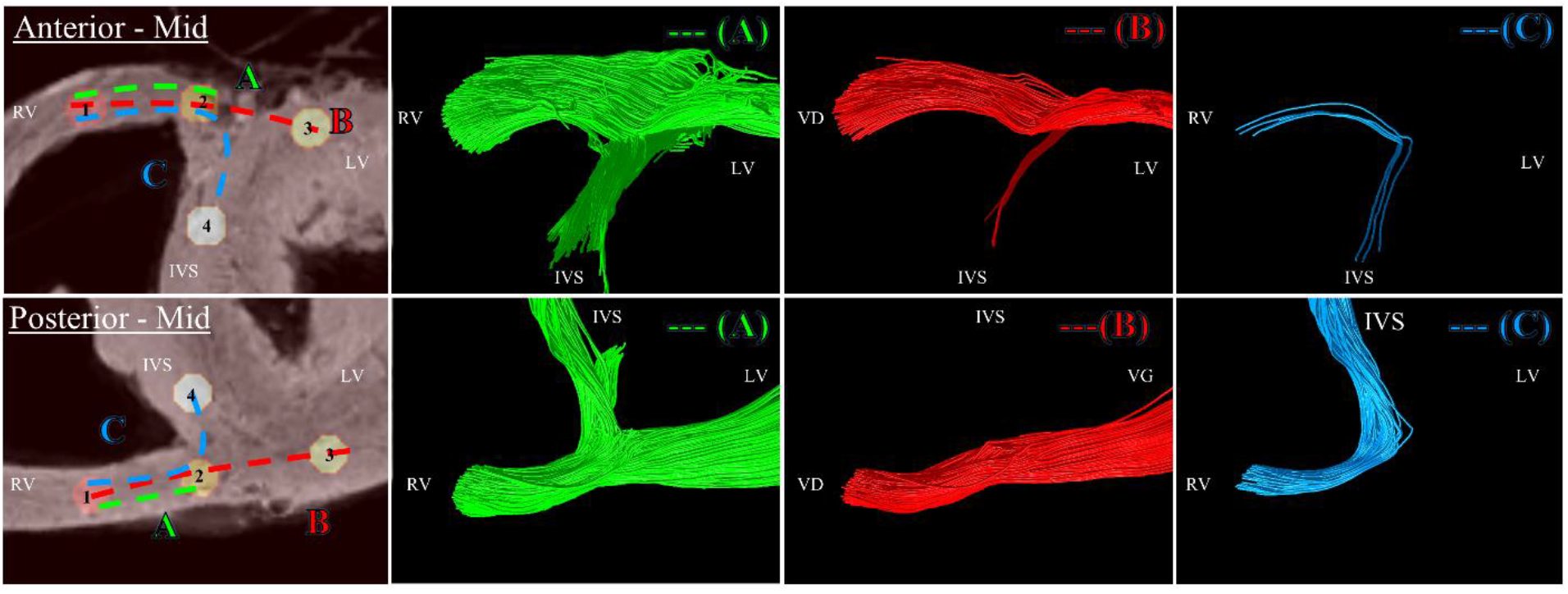
Tractography of the junction obtained with a seeding performed in the mid-ventricular area. Left windows: four cylindrical ROIs: R1 (red), R2 (yellow), R3 (light yellow), R4 (white) were manually defined at the LV/RV junction in the basal area for the anterior (top) and posterior (down) junction. Three arbitrary pathways: A (from ROI R1 to R2) B (from ROI R1 to R2 to R3) C (from ROI R1 to R2 to R4) were also defined as input for computing the streamlines. ROIs and pathways were overlaid on anatomical template images. The other windows correspond to resulting streamlines in 3D view for each pathway. Anterior (Top) Streamlines split into two output tracts. The fascicle of streamlines going to the IVS leaves the image plane and goes to the wall of the pulmonary artery as described in Figure 6. Pathway (B) and (C) were found connecting. Posterior (Down) Streamlines split into two output tracks located in the IVS, in the free wall (A) but do not cross the posterior singularity. Pathways (B) and (C) were found connecting.

Table 1 summarizes the streamline frequency percentages for the 3 samples in apex, mid-ventricular, base areas for the three tracking algorithms. The ratios are all equal to 100 % for pathway A and close to 100% for pathway B except for the posterior basal area (69.2%, 2.1%, 62.3% using the DET algorithm for samples #1,#2,#3 respectively) and the anterior apex area (100%, 32.1%, 2.6% using the DET algorithm for sample #1,#2,#3 respectively). For pathway C, the ratios are close to 0% in the anterior junction (red square) and range 0% from 100% in the posterior junction (green square). The DET and PROB algorithms gave approximately the same ratio, the use of the FACT algorithm tends to decrease the ratio by a factor of 2 or more. Variability over samples is noticeable for pathway B in the posterior mid area and the anterior apex area and for pathway C in the posterior basal and apex area. The corresponding ratio are 100%, 2.4%, 100% and 13.6% 0% 97.1% for sample #1, #2, #3.

**Table 1:**
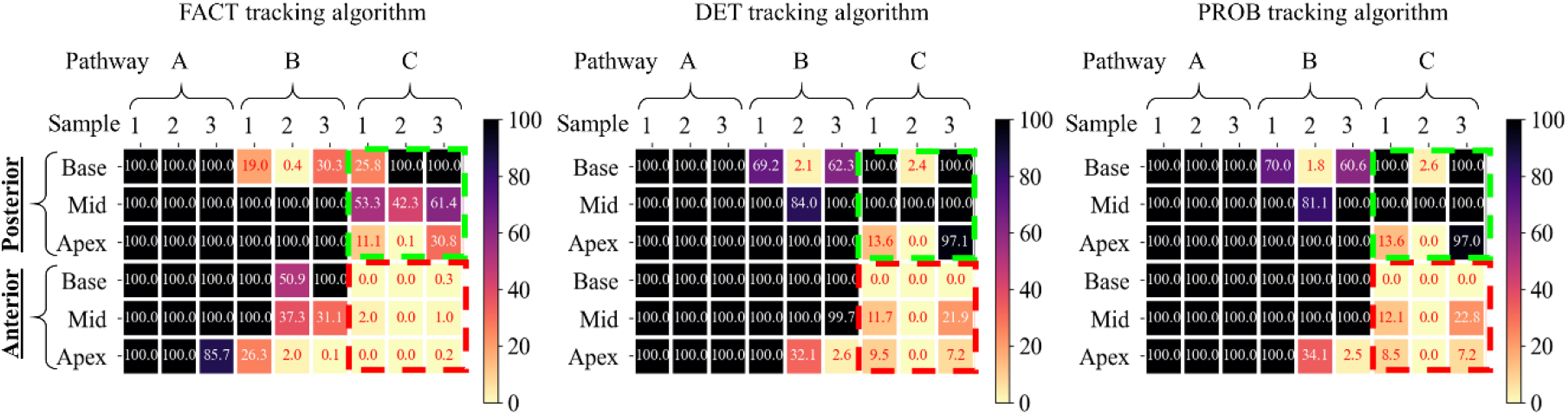
Global myofiber architecture of the junction using seed-based tractography. Influence of the tracking algorithms. Streamlines count as a percentage for the 3 samples in apex, mid-ventricular, base areas for the 3 different tracking algorithms. The ratio is defined as the number of resulting streamlines over the total number of requested streamlines. Variation in connectivity and variability over samples are particulary noticeable for the pathway C. Nevertheless, in all cases, the ratios were found inferior for pathway C for all areas but remained close to 100 % (square with green dotted line) in the posterior junction for the base and mid-ventricular level while values were close to 0 % in the anterior junction (square with red dotted line).

## Discussion

Using anatomical and DW images, we proposed a new framework combining atlas creation and tractography to characterize the cardiac microstructure of *ex-vivo* sheep hearts. We first built a 150 μm isotropic anatomical template and a 300 μm isotropic voxel size FA template. Assessment of the accuracy of inter-subject spatial normalization was done qualitatively by visual inspection. Although misregistration of vessels was visible in different areas of the myocardium, the proposed pyramidal approach to template creation resulted in excellent anatomical consistency.

The main novelties of the proposed template were to cover both ventricles, the atria, and a fraction of the arteries at an unprecedented spatial resolution of 600 μm isotropic for the DTI acquisition. The diffusion metric distributions were found close to the individual sample (Supplementary Fig. 2) and agree with prior studies on other species [36, 40] ensuring that the proposed DTI template creation framework preserves the quantitative DTI metrics. Although image-based quantitative measurements were made to assess the accuracy of the model (Supplementary Fig. 1), the preservation of the FA distribution is also a strong argument. In case of mis-registration, tensor averaging would have decreased the anisotropy [56].

As shown in Fig. 4, spatial variation of the FA was observed in individual and averaged diffusion tensors. This could indicate i) intra-voxel variability with an abrupt change of fiber orientation within the voxel leading to reduced anisotropy (in the mid-wall of the septum) ii) the presence of inter-subject anatomy variability (close the edge of the posterior singularity in the averaged diffusion tensor) iii) the potential existence of registration errors at the voxel scale. A combination of all three scenarios is also likely probable.

The study emphasizes the use of tractography to depict the cardiac fiber organization. This approach is commonly used in neurosciences to study brain connectivity [57–59]. Except for the studies mentioned by Sosnovik et al [60], its use in cardiac diffusion remains marginal and focuses mostly on whole heart tractography for visualization purposes. It is important to highlight that there is an order of magnitude between the size of cardiomyocytes (around 120 μm × 40 μm in diameter [61]) and both the minimum length of the computed streamline (1 mm) and also the voxel size of the reconstructed image (300 × 300 × 300 μm^3^). In this work, the streamlines were used to give an insight of the myofiber architecture, Figures 5–7 show how this 3D representation depicts both the local variation of the fiber orientation but also the continuity of the fiber in the surrounding area. The continuity is dependent on the DW analysis methods, the tracking algorithms and the input parameters [62]. In this study, comparison of diffusion-weighting techniques was not possible due to the unique b-value and the acquisition of 6 diffusion directions. We compared only the use of three different tracking algorithms. The angle parameter of 60° was found to be a maximum to maintain the physical behavior of the fiber (an angle of 90° or more results in undulating streamlines). The cut-off parameter was set to 0.1 to maximize connectivity as shown in Supplementary Fig. 4. The max length parameter was easier to set either depending on the expected size of the object or the desired rendering. Pitfalls of tractography techniques are numerous [63] and have been raised in many studies even with state-of-the-art algorithms [57, 64]. Nevertheless, investigation of the spatial continuity was found difficult using only a visual assessment of the eigenvectors on 2D slices or with a Multiplanar reformation or reconstruction (MPR) viewer.

In our work, the first limitation is the low angular resolution of the DW dataset. A larger number of diffusion directions is recommended for a better estimation of the diffusion tensor [65]. Recent *ex vivo* cardiac studies reported a number of gradient directions from 7 [66], 30 [67], 32 [68] to 61 [66]. Nevertheless, as demonstrated in [66], the trade-off between signal-to-noise ratio and angular resolution are not the only parameters to take into account. The heterogeneity of diffusion directions is better depicted using a high spatial resolution and without voxel anisotropy limiting the voxel averaging of the DTI metrics. By focusing on the spatial resolution and SNR, the proposed approach excludes the acquisition of high angular sampling of the Q-space in regard to the long scan time and prevents from applying more advanced diffusion characterization models like q-ball imaging (QBI) or diffusion spectrum magnetic resonance imaging (DSI).

The second limitation is the tensor model that constrains the fiber orientation to one unique direction per voxel. DSI tractography could provide a more realistic representation of the myofiber architecture [60]. The democratization of undersampled acquisition [28] in pre-clinical settings could also be a potential solution to increase the angular resolution while limiting the scan time.

To demonstrate the added value of the tractography, the structural organization of the anterior and posterior junction was characterized. The study focuses on one fiber-bundle in the posterior junction and three fiber-bundles in the anterior junction. Tractography in a selected ROI was useful to visualize the spatial coverage of the region independently of the surrounding environment (Fig. 5A). Tractography with a specific region seeding (eg. the user manually delineates only a small ROI in the fiber bundle) was useful to display the fiber organization without apriori knowledge of the global myofiber arrangement (Fig. 5C-E, Fig. 7E-H). In the last part of the study (Fig. 8 and Supplementary Fig. 5), generated streamlines were constrained by a user rule and must traverse all specified ROIs to be included in the final tractogram. The ROI was defined on both sides of the junction at three different levels of the ventricle (apex, mid, base). Note that the tested rule did allow crossing pathways between levels and represent a fraction of the structural organization. We then analyzed qualitatively and quantitatively the number of generated streamlines that respect a user rule. A ratio of 100% indicating that all requested streamlines have been generated by the tracking algorithms. Therefore the resulting value is not only dependent on the tracking parameters but also on the request number of streamlines. In all cases, the ratios were found inferior for pathway C for all areas but remained close to 100 % in the posterior junction for the base and mid-ventricular level while values were close to 0 % in the anterior junction indicating the absence of connectivity.

The spatial resolution of 0.6 mm isotropic, and the acceptable inter-variability over subject, enabled the identification of fibers in the IVS and LV that present either a rapid change in orientation or some deviation or a notable deviation from the circular arrangement of the fiber depicted by the helix angle rule. At first glance, we noticed a more complex structural organization of the anterior junction in comparison to the posterior junction. The global myofiber organization was found closely linked to the surrounding structure including the pulmonary and the aorta. The “Y” pattern, depicted in Fig. 7B, is also associated with the location of the anterior-septal perforator arteries recently mentioned in [69]. Finally, visual inspection and streamline connectivity of the anterior junction suggest that most fibers end in the anterior wall while a strong connectivity fiber is visible in the posterior wall. Indeed, a smoother transition of fiber orientation was noticed in the posterior wall (gradient of green to purple in Fig. 4) in the endocardial part of the LV while fiber orientation changes abruptly between adjacent voxels in the IVS in agreement with [70] and the middle of the LV/RV junction (blue in Fig. 4) depicting a triangular shape in the coronal view. We found no reference and possible comparison in the literature regarding the global myofiber organization in this region. The macroscopic size (1 cm^3^ volume) of this substructure suggests that experimental observation of the myofiber orientation could be seen by eye without magnification. We proceeded to a dissection of the heart of another animal that confirmed the presence of this substructure.

The finding must be balanced for two differents reasons: i) the fix state of the ex-vivo samples and ii) the low number of datasets. i) Hearts cardiac muscle is fully relaxed due to cardioplegic flushing before formalin fixation. Therefore, hearts were imaged in relaxed state but were not in a complete diastolic phase. A direct comparison with *in vivo* DTI measures is complicated because the state of *ex vivo* hearts does not matched the *in vivo* contractile state[16] [47]. Therefore, the finding cannot be directly transposed to *in vivo* where the fiber angles vary throughtout the cardiac cycle. However,limited changes are observed for fibers organization [18, 20, 71] between *in vivo* cardiac phase (systolic and diastolic) and *ex vivo* states (relaxed and contracted) and only sheetelets orientations are highly dependent of contractile state. Thus, this fiber-bundle which is aligned in the *ex vivo* state perpendicular to the usual fiber orientation is probably present *in vivo* at any contraction stage. ii) The use of three hearts only is a limitation since it does not allows to explore the inter-subject variability. However, our study focuses on a macroscopic area which present in the three individuals. Moreover, efforts were dedicated to limit the inter subject variability by using animals of identical strain, age and sex as well as by using the exact same sample preparation protocole. As a comparison the first atlas of the human heart was built with 10 hearts [37] while using individal of different ages and sex which could induce important inter-subject variability. That the reason why this number appear sufficient to enforce the conclusion of our study and thus was kept low for ethical consideration.

Finally, mechanical and functional aspects are not discussed in the current study but the individual DW images, diffusion tensors, averaged diffusion tensors of the junction are already available in the Zenodo platform and delivered with shell script for computing the diffusion tensor maps and the streamlines. The conversion of the eigenvectors to CSV format for mesh generation and integration into EP/mechanical models is also possible (see Data availability section) and the visualization of the results can be done with well documented open-source software.

## Conclusion

Demonstrating that the myocardial–myocyte orientation or myolaminar/sheetlet structure changes are correlated with a pathology could theoretically allow the definition of relevant quantitative markers in clinical routine to diagnose heart pathologies. A whole heart diffusion tensor template representative of the global myofiber organization over species is therefore crucial for comparisons across populations. In this paper, we proposed a new framework combining template creation and tractography to characterize the cardiac microstructure of three *ex vivo* sheep hearts and demonstrate the potential application by providing a novel description of the ventricular junction in large mammalian sheep.

## List of abbreviations

MRI: Magnetic resonance imaging
DTI: Diffusion tensor imaging
FLASH: Fast low angle shot
FA: Fractional anisotropy
DWI: Diffusion Weighted imaging
SNR: Signal-to-noise ratio
ADC: Apparent Diffusion Coefficient
cFA: color coded FA
ROI: Region of interest
LV: Left ventricle
RV: Right ventricle
IVS: intraventricular septum
PA: pulmonary artery
AO: aorta
TV: tricuspid valve
HA: Helix angle

## Ethics approval and consent to participate

Animals protocol was approved by the Animal Research Ethics Committee (Comité d’Ethique en Expérimentation Animale de Bordeaux; CEEA50) in accordance with the European rules for animal experimentation (European legislation 2010/63/UE; 2010).

## Consent for publication

Not applicable

## Availability of data and materials

A large fraction of the datasets analyzed during the current study has already been released and more are available upon reasonable request. Data are available at this link (https://doi.org/10.5281/zenodo.5140252). Results are visible with the mrview viewer using the panels overlay/tensor/tractography. Command scripts that computes in native space (diffusion tensor, first eigenvectors, tractography) and command-line instructions for generating figures are also available (https://github.com/valeryozenne/Cardiac-Structure-Database). The remaining fraction of the data is still being analyzed for other purposes and cannot be made publically available at this time.

## Authors’ contributions

VO designed the study. JM performed the acquisition of data. JM, YBP and VO performed the development and data processing. MY and VO performed the dissection study. VO wrote the initial draft of the final paper. JM, MY, YBP, and VO edited the draft and provided important scientific discussion for the finalization of the manuscript. All authors read and approved the final manuscript.

## Acknowledgements

The authors are grateful for the help provided via the Github or Discourse platform by Philip Cook and Nick Tustison regarding the use of ANTs and Max Pietsh, Robert Smith and Donald Tournier regarding the use of MRtrix3 software. The authors thank Girish Ramlugun for proofreading the article.

## Authors’ information (optional)

