## Supplementary figures for "A groupwise registration and tractography framework for cardiac myofiber architecture description by diffusion MRI: an application to the ventricular junctions"

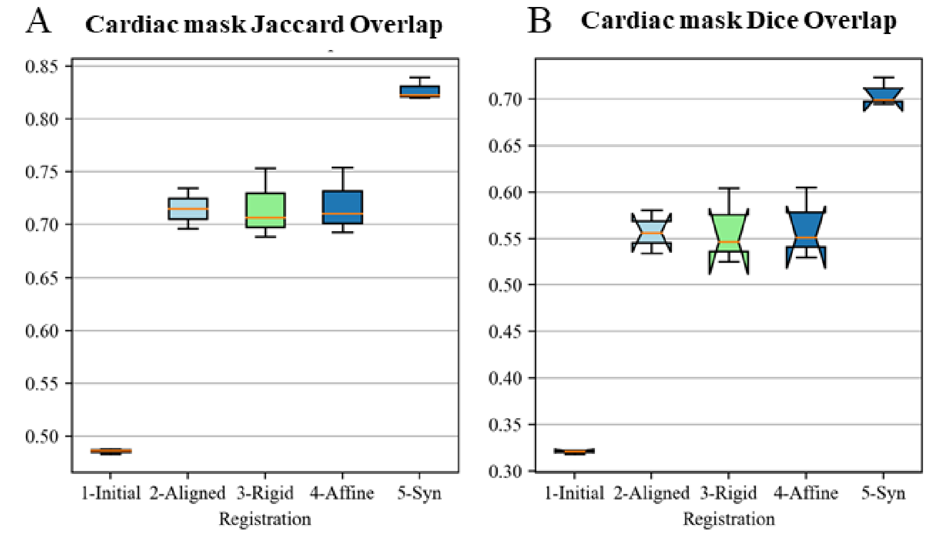


C **P-values for difference in performance measured by paired student's T-test**

| | 1-Initial vs 5-SyN | 2-Aligned vs 5-SyN | 3-Rigid vs 5-SyN | 4-Affine vs 5-SyN |
|---|---|---|---|---|
| Dice | 0,0000006 | 0,0008863 | 0,0052270 | 0,0047301 |
| Jaccard | 0,0000017 | 0,0007581 | 0,0042474 | 0,0038707 |

**Supplementary Figure 1. Comparison of registration quality using Jaccard index and the Dice Similary Coefficient.** The box and whisker plots show the Jaccard (A) and Dice (B) metrics computed between each individual cardiac mask (or label) after different transformation models. Pairwise student T-test was used to determine if performance differences are significant. The Symmetric Normalization (SyN) transformation model showed superior performance over other transformation model.

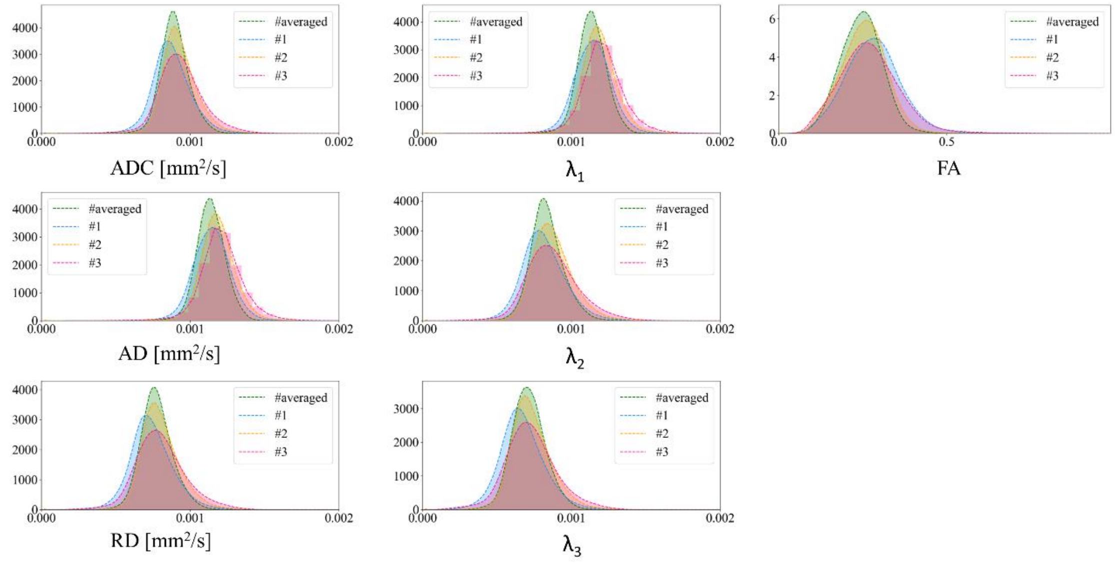


**Supplementary figure 2:** Diffusion metric distributions derived from the individual (#1 blue, 2# yellow, #3 red) and template diffusion tensors (green). The computed metrics are the ADC, the axial diffusivity (AD), the radial diffusivity (RD), the three eigenvalues ($\lambda_1$, $\lambda_2$, $\lambda_3$), and the FA.

**Supplementary figure 3 a, b, c, d, and e: Animated visualization of the cFA maps for the three individual and averaged diffusion tensors.** Movies with cFA maps overlaid an anatomical image for all individual three samples after rigid alignment of the long axis of the left ventricle along the z-axis of the volume. A rotation of 360° by step of 1° is displayed during the video. The same representation is used for the template diffusion tensor. Background and vessels were masked and can be visible as small black dots.

- At 0-1s, in the left view, a fiber arrangement (in blue) is visible from the basal area to the mid-ventricular level close to the papillary muscle.
- At 5-6s, in the left view, a fiber arrangement (in blue) is visible at the intersection of the LV and RV from the basal area to the mid-ventricular level.
- At 9-10 s, in the central view, a fiber arrangement (in blue) is visible in the basal area of the IVS, divided into two bundles of fibers that go on the one hand to the pulmonary artery and on the other hand to the aorta.

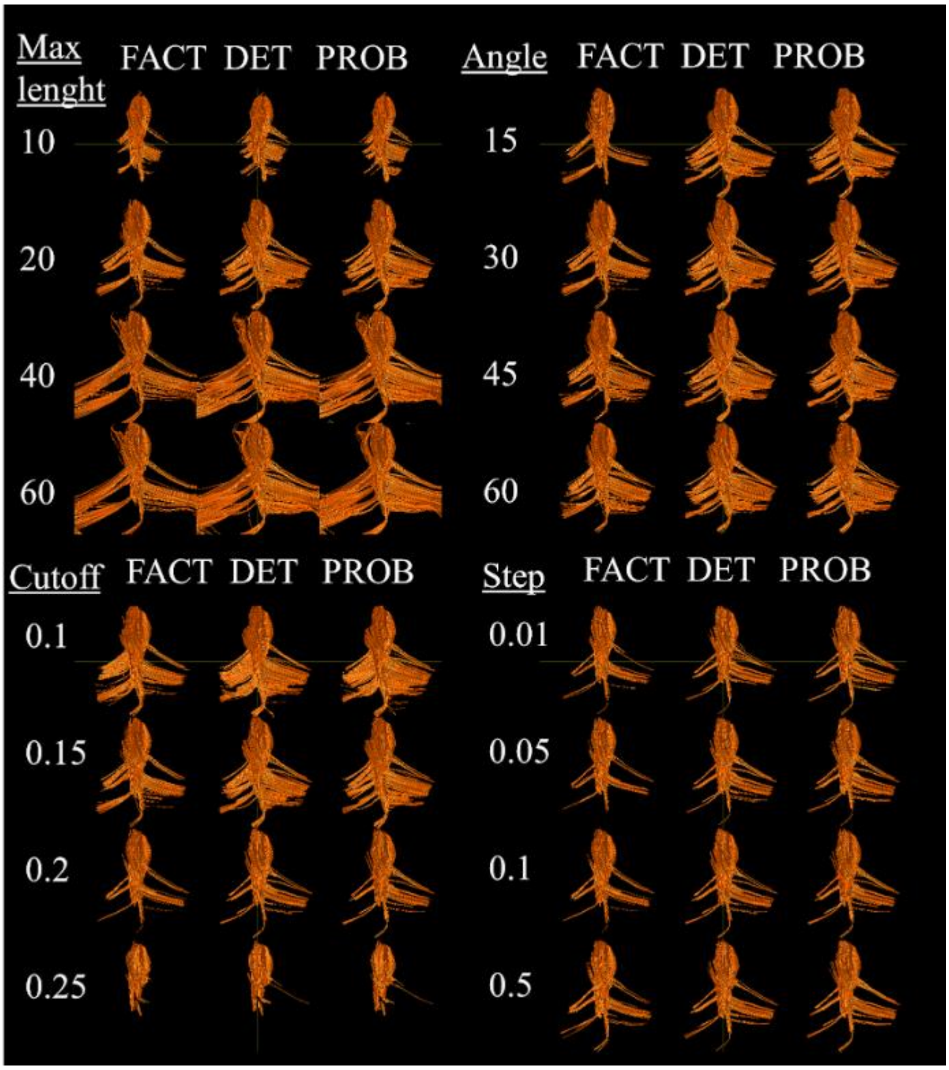

**Supplementary figure 4: Tractography performed in the posterior singularity with different tuning parameters.** All streamlines were obtained for sample #1 in native space using the same seeds and warped in template space to ease the comparison with the previous figure.

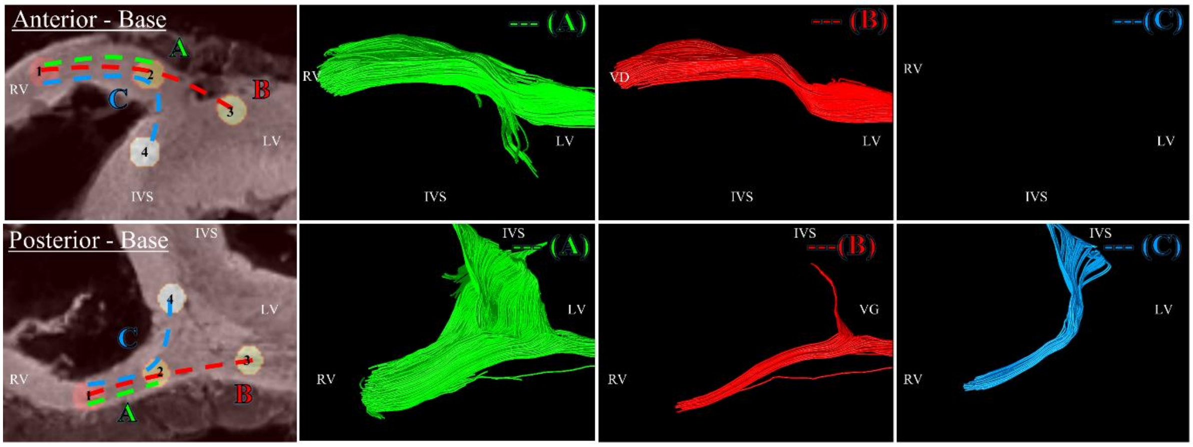


**Supplementary Figure 5: Tractography of the junction obtained with a seeding performed in the basal area.** Left windows: four cylindrical ROIs: R1 (red), R2 (yellow), R3 (light yellow), R4 (white) were manually defined at the LV/RV junction in the basal area for the anterior (top) and posterior (down) junction. Three arbitrary pathways: A (from ROI R1 to R2) B (from ROI R1 to R2 to R3) C (from ROI R1 to R2 to R4) were also defined as input for computing the streamlines. ROIs and pathways were overlaid on anatomical template images. The other windows: resulting streamlines in 3D view for each pathway. Anterior (Top) Streamlines split into two output tracts. The pathway (C) was not found connecting. Posterior (Down) Streamlines split into multiple output tracts located either in the IVS, at the junction, or in the free wall (A). Existing pathways (B) and (C) were found.
